# Selective prefrontal-amygdala circuit interactions underlie social and nonsocial valuation in rhesus macaques

**DOI:** 10.1101/2021.04.13.439737

**Authors:** Maia S. Pujara, Nicole K. Ciesinski, Joseph F. Reyelts, Sarah E.V. Rhodes, Elisabeth A. Murray

**Author notes:** **Corresponding author:** Elisabeth A. Murray, Ph.D., Laboratory of Neuropsychology, NIMH, Building 49, Suite 1B80, 49 Convent Drive, Bethesda, MD 20892-4415 USA. **Financial Interests/Conflict of Interest:** The authors declare no competing financial interests.

## Abstract

Lesion studies in macaques suggest dissociable functions of the orbitofrontal cortex (OFC) and medial frontal cortex (MFC), with OFC being essential for goal-directed decision making and MFC supporting social cognition. Bilateral amygdala damage results in impairments in both of these domains. There are extensive reciprocal connections between these prefrontal areas and the amygdala; however, it is not known whether the dissociable roles of OFC and MFC depend on functional interactions with the amygdala. To test this possibility, we compared the performance of male rhesus macaques (*Macaca mulatta*) with crossed surgical disconnection of the amygdala and either MFC (MFC x AMY, *n*=4) or OFC (OFC x AMY, *n*=4) to a group of unoperated controls (CON, *n*=5). All monkeys were assessed for their performance on two tasks to measure: (1) food-retrieval latencies while viewing videos of social and nonsocial stimuli in a test of social interest, and (2) object choices based on current food value using reinforcer devaluation in a test of goal-directed decision making. Compared to the CON group, the MFC x AMY group, but not the OFC x AMY group, showed significantly reduced food-retrieval latencies while viewing videos of conspecifics, indicating reduced social valuation and/or interest. By contrast, on the devaluation task, group OFC x AMY, but not group MFC x AMY, displayed deficits on object choices following changes in food value. These data indicate that the MFC and OFC must functionally interact with the amygdala to support normative social and nonsocial valuation, respectively.

**Significance Statement:** Ascribing value to conspecifics (social) vs. objects (nonsocial) may be supported by distinct but overlapping brain networks. Here we test whether two nonoverlapping regions of the prefrontal cortex, the medial frontal cortex and the orbitofrontal cortex, must causally interact with the amygdala to sustain social valuation and goal-directed decision making, respectively. We found that these prefrontal-amygdala circuits are functionally dissociable, lending support for the idea that medial frontal and orbital frontal cortex make independent contributions to cognitive appraisals of the environment. These data provide a neural framework for distinct value assignment processes and may enhance our understanding of the cognitive deficits observed following brain injury or in the development of mental health disorders.

## Introduction

Both individual and collective fitness contribute to the evolutionary success of anthropoid primates, including macaques. In their natural habitats, macaques gain advantages for survival by optimizing behaviors that lead to beneficial outcomes both for themselves and their group. In a laboratory setting, monkeys, like humans, develop and express transitive subjective preferences for different foodstuffs (Tremblay and Schultz, 1999; Padoa-Schioppa et al., 2006; Martin et al., 2018) and juices (Tremblay and Schultz, 1999; Padoa-Schioppa and Assad, 2006). Macaques also forego juice rewards to view images of female perinea (i.e., sex skin) and dominant male monkey faces (Deaner and Platt, 2003; Deaner et al., 2005), which suggests that social cues have intrinsic value. Although assigning value to foods and/or conspecifics and the stimuli that predict them is a central feature of cognition, evidence suggests that these processes are supported by distinct neural networks. In this study, we examined the neuroanatomical basis of two aspects of biological value in macaques: nonsocial objects associated with food rewards and social signals conveyed by conspecifics.

Neuropsychological research in humans points to a causal role for the amygdala and ventromedial frontal cortex (VMF) in carrying out these essential higher-order functions. Patients with Urbach-Wiethe disease who experience selective bilateral amygdala damage are profoundly impaired both in their ability to learn from social information (Rosenberger et al., 2019) and making rational decisions (Brand et al., 2007; Talmi et al., 2010). Similarly, patients with lesions of the VMF, which typically compromises substantial portions of both orbital and medial sectors of the frontal cortex, are impaired in both social cognition (Eslinger and Damasio, 1985; Barrash et al., 2000; Hornak et al., 2003; Mah et al., 2004; Beer et al., 2006; Ciaramelli et al., 2007; Jenkins et al., 2014; Hiser and Koenigs, 2018) and rational or goal-directed decision making (Eslinger and Damasio, 1985; Anderson et al., 1999; Fellows and Farah, 2003; Camille et al., 2004; Koenigs and Tranel, 2007; Wheeler and Fellows, 2008; Pujara et al., 2015; Reber et al., 2017; Barrash et al., 2018; Hiser and Koenigs, 2018).

Because brain damage in humans rarely respects boundaries of individual structures, teasing apart the contributions of the medial frontal cortex (MFC) and orbitofrontal cortex (OFC) and their independent networks depends on studies in nonhuman primates, where targeting these areas individually and testing circuit interactions is more tractable (Vaidya et al., 2019). Studies in macaques suggest specializations for MFC and OFC in social and nonsocial valuation, respectively. Whereas the OFC is involved in value coding, updating stimulus-outcome associations, and value-based decision making (Padoa-Schioppa and Assad, 2006; Padoa-Schioppa, 2011; Wallis, 2011; Murray and Rudebeck, 2018), the MFC has been implicated in social cognition (Rudebeck et al., 2006; Noonan et al., 2010a; Lockwood et al., 2020) and encoding of rewards for the self and others (Chang et al., 2013). Given the known reciprocal anatomical connections between the amygdala and both the MFC and OFC (Carmichael and Price, 1995; Ghashghaei and Barbas, 2002; Saleem et al., 2008), it is possible that the MFC and OFC must functionally interact with the amygdala to support these processes (Gangopadhyay et al., 2021). If so, these prefrontal-amygdala circuits may be separable in empirical tests of their functions.

Although there is evidence that OFC-amygdala interactions support value-based decision making by Baxter et al. (2000), the aspiration OFC lesions in that study may have involved damage to fibers of passage (Rudebeck et al., 2013b); therefore, using selective, excitotoxic lesions to update the interpretation of this circuit’s function would be timely. No macaque lesion studies to date have examined OFC- and MFC-amygdala interactions in social cognition. We predicted that, relative to intact controls, monkeys with crossed surgical disconnection of the MFC and the amygdala would be impaired in evaluating naturalistic videos of conspecifics, but not goal-directed decision-making, whereas monkeys with crossed surgical disconnection of the OFC and the amygdala would show the opposite pattern of results.

## Materials and Methods

### Subjects

Thirteen adult male rhesus monkeys (*Macaca mulatta*) served as subjects. Four monkeys sustained unilateral aspiration lesions of the MFC targeting area 32 (Carmichael and Price, 1994) and four monkeys sustained unilateral excitotoxic lesions of the OFC targeting Walker’s areas 11, 13, and 14 (Walker, 1940). All operated monkeys sustained excitotoxic amygdala lesions in the hemisphere contralateral to their cortical lesions (i.e., crossed lesions). The remaining five monkeys were retained as unoperated controls (CON). All training and testing was conducted postoperatively, starting with an action-outcome reinforcer devaluation task previously described in Fiuzat et al. (2017), followed by a social valuation task and an object-outcome reinforcer devaluation task described below. Importantly, all monkeys had the same training history.

Monkeys weighed between 7.1 kg and 15.2 kg (mean: 9.5 kg) and all were at least five years old at the start of testing. Each animal was housed individually with visual and auditory access to conspecifics in the animal housing room, kept on a 12-h light-dark cycle (lights on at 7:00 A.M.), maintained on controlled access to primate chow supplemented with fruit to ensure sufficient motivation to respond in the test apparatus, and given access to water 24 hours a day. Testing occurred during the light period. All procedures were reviewed and approved by the NIMH Animal Care and Use Committee.

### Surgery

Eight monkeys received surgery in two stages to produce crossed surgical disconnection of the amygdala and either the MFC or OFC. All operated monkeys sustained amygdala damage in one hemisphere and damage of either MFC (MFC x AMY) or OFC (OFC x AMY) in the other hemisphere; groups were balanced for site (frontal cortex, amygdala) and hemisphere (left, right) of first surgery. For the purpose of relating the location of our intended lesions to other commonly employed anatomical frameworks, we note that the intended MFC lesion in our study corresponds approximately to area 32, also known as prelimbic cortex or pregenual cortex (Barbas and Pandya, 1989; Carmichael and Price, 1994), and the intended OFC lesion corresponds approximately to areas 11l, 11m, 13l, 13m, 13b, 14r, 14c, and 10m (Carmichael and Price, 1994). The intended amygdala lesion encompassed both the basolateral and centromedial nuclear groups.

At the time of surgery, anesthesia was induced with ketamine hydrochloride (10 mg/kg, i.m.) and maintained with isoflurane (1.0-3.0%, to effect). Heart rate, respiration rate, blood pressure, expired CO_2_, and body temperature were monitored during surgery and isotonic fluids were given throughout. Aseptic procedures were used. After completing the series of ibotenate injections (excitotoxic) or combination of suction and cautery (aspiration), the surgical site was closed in anatomical layers with sutures. The preoperative and postoperative treatment regimen consisted of dexamethasone sodium phosphate (0.4 mg/kg, i.m.) and cefazolin antibiotic (15 mg/kg, i.m.) for one day before surgery and one week after surgery to reduce swelling and prevent infection, respectively. At the end of surgery and for two additional days, the monkeys received the analgesic ketoprofen (10-15 mg, i.m.), followed by ibuprofen (100 mg) for the following five days. Operations were separated by a minimum of two weeks. Postoperative behavioral testing was initiated 10-14 days following the second stage of surgery.

Monkeys receiving an amygdala lesion were anesthetized and then placed in a stereotaxic frame. A large bone flap was turned over the appropriate portion of the cranium. The injection sites were calculated based on landmarks that were visible in the MRI scans obtained before surgery. The sagittal sinus served as a landmark for the mediolateral coordinates and the interaural plane (ear bars) served as a landmark for the anteroposterior and dorsoventral coordinates. The monkeys received between 16 and 23 unilateral injections to sites located ∼2 mm apart in each plane. Each injection consisted of 1.0 *μ*l of ibotenate (10-15 *μ*g/ul; 0.2 *μ*l/min; Sigma-Aldrich) administered via a 30-gauge Hamilton syringe held in a David Kopf Instruments manipulator. Before lowering the needle, small slits were made in the dura to allow the needle to pass unobstructed into the brain. The needle remained in place 2-3 min after each injection to limit diffusion of the toxin up the needle track. On closing, 30 ml of mannitol (25%, 1 ml/min, i.v.) was administered to control edema.

Monkeys receiving either MFC or OFC lesions were anesthetized and then placed in a custom head holder. At the beginning of surgery, monkeys were given 30 ml of mannitol (25%, 1 ml/min, i.v.) to increase access to the orbital and medial surfaces and to control edema. For the MFC lesion, a large, bilaterally symmetrical bone flap was turned over the frontal cortex, and the dura was reflected toward the midline. Sulcal landmarks on the medial surface were identified with the aid of an operating microscope, and the boundaries of the lesion were marked with a line of electrocautery. Then, using a combination of suction and electrocautery, the MFC was removed by subpial aspiration through a fine-gauge metal sucker that was insulated except at the tip. The intended MFC lesion was approximately rectangular in shape and extended rostrocaudally along the length of the rostral sulcus from the genu of the corpus callosum, caudally, to the rostral tip of the cingulate sulcus, rostrally. The dorsal boundary of the lesion was ∼2 mm ventral to the cingulate sulcus (at the midlevel of the genu of the corpus callosum), and the ventral boundary of the lesion was the fundus of the rostral sulcus.

For the OFC lesion, a large bone flap was turned over the dorsal frontal cortex. The dura was opened with a crescent-shaped cut and then reflected toward the orbit. Sulcal landmarks on the orbital surface were identified with the aid of an operating microscope. Unilateral injections of ibotenate were made into sites located ∼2 mm apart. At each site, 1.0 *μ*l of ibotenate (10-15 *μ*g/*μ*l; Sigma-Aldrich) was injected as a bolus via a hand-held Hamilton syringe with a 30-gauge needle. Monkeys received between 87 and 99 injections. The intended OFC lesion extended from the fundus of the lateral orbital sulcus, laterally, to the rostral sulcus on the medial surface of the hemisphere, medially. The rostral boundary of the intended lesion was an imaginary line joining the rostral tips of the medial and lateral orbital sulci. The caudal boundary of the intended lesion was an imaginary line joining the most caudal points of the medial and lateral orbital sulci.

### Lesion Assessment

For the OFC and the amygdala excitotoxic lesions, the estimated damage for each monkey was assessed using a T2-weighted MRI scan obtained 4-8 days after surgery. The location and extent of excitotoxic lesions in the hippocampus is reliably indicated by the white hypersignal present on the T2-weighted postoperative scans (Malkova et al., 2001). For the amygdala and surrounding structures, however, the hypersignal may represent an overestimation of damage (Basile et al., 2017). For the MFC aspiration lesions, the estimated damage for each monkey was assessed using a T1-weighted scan obtained postoperatively. Two of the four monkeys with MFC aspiration lesions (Cases #3 and #4) had scans that were acquired approximately three to four months after surgery, whereas the remaining two (Cases #1 and #2) had scans that were acquired five to seven years after the surgery.

We developed a semi-automated lesion mapping procedure to assess the location and extent of the lesions. For all T2-weighted scans, volume estimates were taken by performing rigid (6-parameter rigid body transformation) and affine (diffeomorphic – allowing for 12-parameter local warps in structure) warps on each scan to the <u>N</u>IMH <u>M</u>acaque <u>T</u>emplate version 2.0 (NMT; 0.5 mm^3^ resolution) (Seidlitz et al., 2018; Jung et al., 2020) using AFNI’s 3dAllineate function for the rigid transform (Cox, 1996; Saad et al., 2009) and antsRegistrationSyN in ANTs for the affine transform (Avants et al., 2011). We then applied thresholding to identify the area of hyperintensity on the transformed T2-weighted scans to generate a binary mask that corresponded to the area of damage. These masks were visually inspected and manually edited to ensure that they fully captured the areas of hyperintensity.

For the T1 scans showing MFC damage, we used AFNI’s @animal_warper (Saad et al., 2009; Jung et al., 2020) to perform a rigid and affine alignment for each subject to the standard NMT version 2.0. For MFC damage, because the aspiration lesion resulted in collapse of nearby tissue into the space created by the lesion, we interpreted the extent of the damage in the transformed T1 scans by comparing the area of damage with the comparable region in the intact contralateral hemisphere, using anatomical and sulcal landmarks of the NMT as references. This method therefore ensured a more accurate estimation of damage than drawing the boundaries of the hypointense areas on the lesioned cortical surface.

A lesion overlap map for each region of interest, distinguished by group for the amygdala, was created by summing the binary masks for each hemisphere and displaying the outputs on the NMT version 2.0 (OFC x AMY, **Fig. 1A**; MFC x AMY, **Fig. 1B**). For each subject, the extent of the damage was estimated by taking the percentage overlap of the lesion mask with the intended lesion mask for each region of interest (MFC, OFC, and amygdala) in its respective hemisphere (OFC x AMY, **Table 1;** MFC x AMY, **Table 2)**.

**Table 1.** Percentage Estimated Damage: OFC x AMY Group. OFC x AMY cases 1-4. Monkeys with injections of ibotenic acid targeting Walker’s areas 11, 13, and 14 unilaterally and injections of ibotenic acid targeting the amygdala in the contralateral hemisphere. Mean: average of the estimated damage to each region.

|  | **OFC**<br>**Lesion**<br>**Side** | **% OFC**<br>**damage** | **Amygdala**<br>**Lesion**<br>**Side** | **%**<br>**Amygdala**<br>**Damage** |
| --- | --- | --- | --- | --- |
| Case 1 | R | 67 | L | 90 |
| Case 2 | L | 78 | R | 84 |
| Case 3 | R | 68 | L | 94 |
| Case 4 | L | 91 | R | 88 |
| **Mean** |  | **76** |  | **89** |

**Table 2.** Percentage Estimated Damage: MFC x AMY Group. MFC x AMY cases 1-4. Monkeys with aspiration lesions approximately targeting Walker’s area 32 unilaterally and injections of ibotenic acid targeting the amygdala in the contralateral hemisphere. Mean: average of the estimated damage to each region.

|  | **MFC**<br>**Lesion**<br>**Side** | **% MFC**<br>**damage** | **Amygdala**<br>**Lesion**<br>**Side** | **%**<br>**Amygdala**<br>**Damage** |
| --- | --- | --- | --- | --- |
| Case 1 | L | 88 | R | 97 |
| Case 2 | R | 89 | L | 97 |
| Case 3 | L | 96 | R | 96 |
| Case 4 | R | 62 | L | 97 |
| **Mean** |  | **84** |  | **97** |

**Figure 1.**
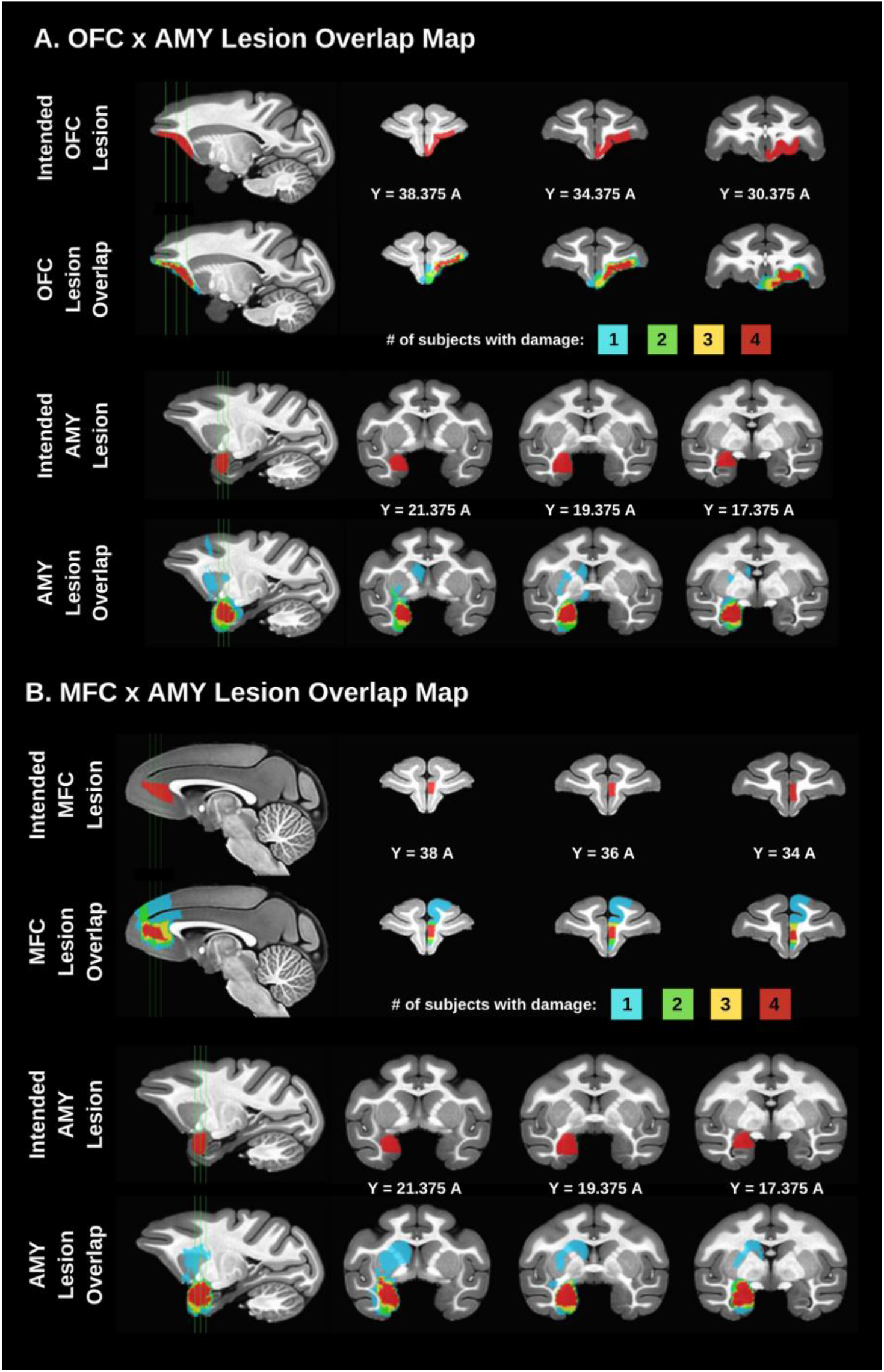
Lesion overlap maps. **A**, damage sustained by the four subjects in the orbitofrontal cortex-amygdala (OFC x AMY) crossed lesion group. **B**, damage sustained by the four subjects in the medial frontal cortex-amygdala (MFC x AMY) crossed lesion group. For visualization purposes, cortical lesions are displayed on the right hemisphere and amygdala lesions are displayed on the left hemisphere. The top row in each panel shows four sections of the intended cortical lesion mask (one sagittal and three coronal views) with corresponding views of the extent of cortical lesion overlap, and the bottom row in each panel shows four sections of the intended amygdala lesion mask (one sagittal and three coronal views) with corresponding views of the extent of amygdala lesion overlap. All coordinates for the NMT version 2.0 are displayed below each section for reference.

In general, the lesions were as intended. Estimated damage to the amygdala ranged from 84% to 97%. Estimated OFC damage ranged from 67% to 91%. Estimated damage to the MFC ranged from 62% to 96%. One monkey in the OFC x AMY group (Case #4) and another monkey in the MFC x AMY group (Case #1) sustained inadvertent damage to the head of the caudate nucleus and putamen, apparently due to an infarction associated with the amygdala injections. Of note, the lesion extents reported for the OFC x AMY group in this report differ from the values reported in Fiuzat et al. (2017) due to a difference in the methodology used to estimate damage.

### Experiment 1: Social Valuation Task

#### Apparatus

Briefly, monkeys were trained in a modified Wisconsin General Testing Apparatus (WGTA) located in a darkened room. Monkeys occupied a wheeled transport cage in the animal compartment of the WGTA. A clear Plexiglas box measuring 11.4 cm (width) x 71.1 cm (length) x 11.4 cm (height) with a hinged back was placed in front of the transport cage in the test compartment of the WGTA. This is the same apparatus used in other studies from this laboratory that have examined responses to objects placed inside the Plexiglas box (Pujara et al., 2019). In this study, however, we modified the apparatus to allow for the presentation of videographic materials, as in Rudebeck et al (2006). Behind the Plexiglas box, a monitor was placed facing the front of the transport cage. On each trial, stimuli presented on the screen were 30-second video clips of one of four social contexts: a large monkey staring; a monkey exhibiting affiliative behaviors (lip-smacking); a small monkey with food in its cheek pouches; and a female macaque with prominent perineal swelling. For further comparison, we used either moving or static neutral images (i.e., objects, scenes) and non-neutral (i.e., emotionally-provocative) predator videos of a slithering snake and a moving toy crocodile. A camera (camera #1) mounted on top of the WGTA recorded each monkey’s movements throughout the trial from a top-down view to track monkeys’ latencies to take food rewards placed on top of the Plexiglas box. Another camera (camera #2) was mounted on top of the monitor and faced the transport cage to record gaze information for subsequent analyses of visual attention to the videos.

#### Procedure

All monkeys were first required to retrieve food rewards that were located on top of the Plexiglas box while the box was empty for 20 trials. By the end of pretraining, monkeys were familiar with the Plexiglas box and the procedure of reaching for rewards overlaying the box. Stimuli were shown in a fixed presentation across two separate test sessions for each monkey. The first test included presentations of the lipsmacking monkey, the female perinea, and the snake videos among seven neutral stimuli for a total of 10 trials. The second test included presentations of the small monkey, the staring monkey, and the crocodile videos among seven neutral stimuli for a total of 10 trials.

On each trial, the opaque moveable screen separating the animal compartment and the test compartment of the WGTA was raised to reveal the video playing on the monitor and a piece of food on the Plexiglas box simultaneously. Therefore, valuation of the content of the video was pitted against the incentive value of the food. Monkeys were given 30 seconds to reach for the piece of food and view the video. After 30 seconds, the screen was lowered to signify the end of a trial. Trials were separated by 30 seconds. Social valuation was operationalized as the difference in the latency to reach for the food reward in the presence of the social videos compared to the neutral stimuli (see **Statistical Analysis** section for the method used to code food-retrieval latencies).

### Experiment 2: Reinforcer Devaluation Task

#### Apparatus

As in Experiment 1, all testing was conducted in the WGTA environment and monkeys were tested inside a wheeled transport cage in the animal compartment of the WGTA. The test compartment of the WGTA held a test tray, which contained two food wells spaced 235 mm apart. Test material for reinforcer devaluation consisted of 120 objects that varied in size, shape, color, and texture. Food rewards for the devaluation task consisted of two of the following six foods: M&Ms (Mars Candies, Hackettstown, NJ), half peanuts, raisins, craisins (Ocean Spray, Lakeville-Middleboro, MA), banana-flavored pellets (Noyes, Lancaster, NH) and fruit snacks (Giant Foods, Landover, MD).

#### Procedure

Procedures for the reinforcer devaluation task have been previously described (Baxter et al., 2000; Rudebeck et al., 2017a). First, each monkey’s preference for six different foods was assessed over a 15-day period. Each day, monkeys received 30 trials consisting of pairwise presentations of the six different foods, one each in the left and right wells of the test tray (e.g., half peanut on the left vs. craisin on the right). The left-right position of the foods was counterbalanced. Preferences were determined by analyzing choices within each of the 15 possible pairs of foods over the final five days of testing. Two foods that the monkey valued highly and that were roughly equally palatable as judged by choices in the food preference test were selected for subsequent behavioral shaping and testing.

Object discrimination learning was employed to set up unique object-outcome associations, followed by reinforcer devaluation tests, in which probe trials gauged monkeys’ abilities to link objects with current food value. For object discrimination, monkeys were trained to discriminate 60 pairs of novel objects. For each pair, one object was randomly designated as the positive object (S+, rewarded) and the other was designated as negative (S-, unrewarded). Half of the positive objects were baited with food 1, and the other half were baited with food 2. For each monkey, the identity of foods 1 and 2 was based on the monkey’s previously determined food preferences. On each trial of discrimination training, monkeys were presented with a pair of objects, each covering a food well, and were allowed to displace one of the two objects. If they displaced the S+ object, they were allowed to retrieve the food, which led to termination of the trial. If they displaced the S-object, no food was available and the trial was terminated. The left-right position of the S+ followed a pseudorandom order. Training continued until monkeys attained the criterion of a mean of 90% correct responses over five consecutive days (i.e., 270 correct responses or greater in 300 trials).

Monkeys’ object choices were then assessed under two conditions for the reinforcer devaluation task: (1) under normal (baseline) conditions, and (2) after one of the foods was devalued via selective satiety. On separate days, four test sessions were conducted, each consisting of 30 trials. Only the positive (S+) objects were used, such that on each trial, a food-1 object and a food-2 object were presented together for choice. Each object covered a well baited with the corresponding food. The object pairs were generated randomly for each session, with the constraint that a food-1 object was always paired with a food-2 object. Preceding two of the test sessions, a selective satiation procedure, intended to diminish the value of one of the foods, was conducted. For the other two test sessions, which provided baseline scores, monkeys were not sated on either food before being tested. The order in which the test sessions occurred was the same for all monkeys and was as follows: (1) baseline test 1; (2) food 1 devalued by selective satiation prior to test session; (3) baseline test 2; (4) food 2 devalued by selective satiation prior to test session. A second test of object choice, identical to the first, was conducted between three to four months after the first object choice test. Monkeys were retrained on the same 60 object pairs to the same criterion as before. After relearning, the object choice test was conducted in the same manner as the first test.

For the selective satiation procedure, a food hopper was filled with a pre-weighed quantity of either food 1 or food 2 and was attached to the front of the monkey’s home cage. The monkey was given a total of 30 minutes to consume as much of the food as it wanted, at which point the experimenter began to observe the monkey’s behavior. Additional food was provided if necessary. The selective satiation procedure was deemed to be complete when the monkey refrained from retrieving food from the hopper for 5 minutes. The amount of time taken in the selective satiation procedure and the total amount of food consumed by each monkey was noted. The monkey was then taken to the WGTA within 10 minutes and the test session was conducted.

We assessed the effect of selective satiation on monkeys’ choices of the foods alone between 30 to 60 days after the second test of object choice. This “food only” test was conducted to evaluate whether satiety transferred from the home cage to the WGTA, and whether behavioral effects of the lesion (if any) were due to an inability to link objects with food value as opposed to an inability to discriminate the foods. This test was identical to both reinforcer devaluation tests 1 and 2, with the important difference that no objects were presented over the two wells where foods were placed. On each trial of the 30-trial sessions, monkeys could see the two foods and were allowed to choose between them. As was the case for object choice tests 1 and 2, there were four critical test sessions, in which two were preceded by selective satiation and two were not.

### Statistical Analysis

#### Social Valuation

Because of the small sample sizes of our experimental groups, we used nonparametric tests for all of our within-test analyses. Food-retrieval latencies were derived from analyses of the video recordings from camera #1, which provided a top-down view of the compartment. Latencies were scored to the nearest frame and had a resolution of ∼4 ms. Time for the latency measure was initiated when the opaque screen was raised ∼15 cm above the test tray. This could be discerned in the videotape by a mark on the cage, visible in the view of camera #2, which provided a frontal view of the monkeys’ behavioral responses to the videos presented on the monitor. The response was considered complete when the monkey grasped the food reward just prior to withdrawing its arm. If no response was made within the trial limit of 30 seconds, a score of 30 seconds was recorded. To observe group differences in duration of time spent viewing the videos, we used the front view from camera #2 to measure the length of time the monkeys looked at the monitor screen that was used to display the stimuli (“look-at duration,” in seconds) for the full 30 seconds of the trial. All measurements were taken by an observer who was naïve to group assignment.

For each monkey, food-retrieval latencies were averaged across both tests for stimuli in the neutral category, nonsocial affective category, and social category. We first tested whether social videos elicited longer looking times compared to neutral stimuli in controls, as shown previously (Rudebeck et al., 2006; Noonan et al., 2010a), by running a within-subject signed-rank Wilcoxon test. To test the effects of crossed surgical disconnections of either the MFC or OFC with the amygdala on social valuation, we then ran separate two-tailed Kruskal-Wallis tests for the social and neutral stimulus categories on the latencies to reach for a food reward, followed by *post hoc* Mann-Whitney *U* tests to compare the experimental lesion groups to the intact control group and to each other.

As a control, to determine whether the groups differed in the amount of time spent viewing the videos, we ran separate two-tailed Kruskal-Wallis tests for the social and neutral stimulus categories on the stimulus look-at durations, followed by *post hoc* Mann-Whitney *U* tests to compare the experimental lesion groups to each other and to the intact control group. Finally, we compared the two lesion groups and controls on their food-retrieval latencies for the nonsocial predator videos (i.e., snake and alligator) to test whether the lesion groups showed altered responses to this category of stimuli.

#### Reinforcer Devaluation

Performance on reinforcer devaluation was measured by calculating a proportion shifted score (Murray et al., 2015) for object choice test 1, object choice test 2, and the food choice test. The proportion shifted score is defined as a shift in the number of food-1 and food-2 associated objects (object test) or food 1 and food 2 (food only test) choices after selective satiation relative to baseline choices, combined across probe tests for food 1 and food 2. The proportion shifted scores obtained from object choice test 1 and object choice test 2 were averaged together to obtain a final object test “proportion shifted” score for each subject. A higher proportion shifted score indicates a greater shift away from choices associated with the devalued food.

To test the effects of crossed surgical disconnections of either the MFC or OFC with the amygdala on value updating, we ran separate two-tailed Kruskal-Wallis tests for proportion shifted scores obtained from object choices, followed by *post hoc* Mann-Whitney *U* test to compare the experimental lesion groups to the intact control group and to each other. As a control, to demonstrate that the groups did not differ in their ability to update choices for the foods themselves following selective satiation (i.e., that impairments were specific to updating stimulus-outcome associations), we ran a separate two-tailed Kruskal-Wallis test for proportion shifted scores for food choices.

#### Double Dissociation

To determine whether there was a double dissociation between the lesion groups, we normalized (*z*-scored) the food-retrieval latencies from the social valuation task (social condition minus neutral condition) and the average proportion shifted scores from the object choice tests. We then ran a 2×2 ANOVA to test for a significant group (OFC x AMY, MFC x AMY) by task (social, nonsocial) interaction, which would indicate a double dissociation between the two groups on task performance.

## Results

### Social valuation

We first evaluated whether social videos attracted the interest of unoperated control monkeys, as shown previously (Rudebeck et al., 2006), indexed by longer food-retrieval latencies in the presence of the social videos compared to neutral stimuli. We found that controls were, on average, slower to reach for food rewards when presented with social (**Fig. 2B**) compared to neutral stimuli (Wilcoxon signed-rank test: *V* = 0, *p* = 0.06; **Fig. 2A**). The group difference approached but did not reach significance due to one control subject who had longer food-retrieval latencies than the other controls for the neutral stimuli (22.25 seconds, greater than one standard deviation above the mean). Thus, in line with previous work, social videos slowed food reward retrieval within the unoperated control group, on average.

**Figure 2.**
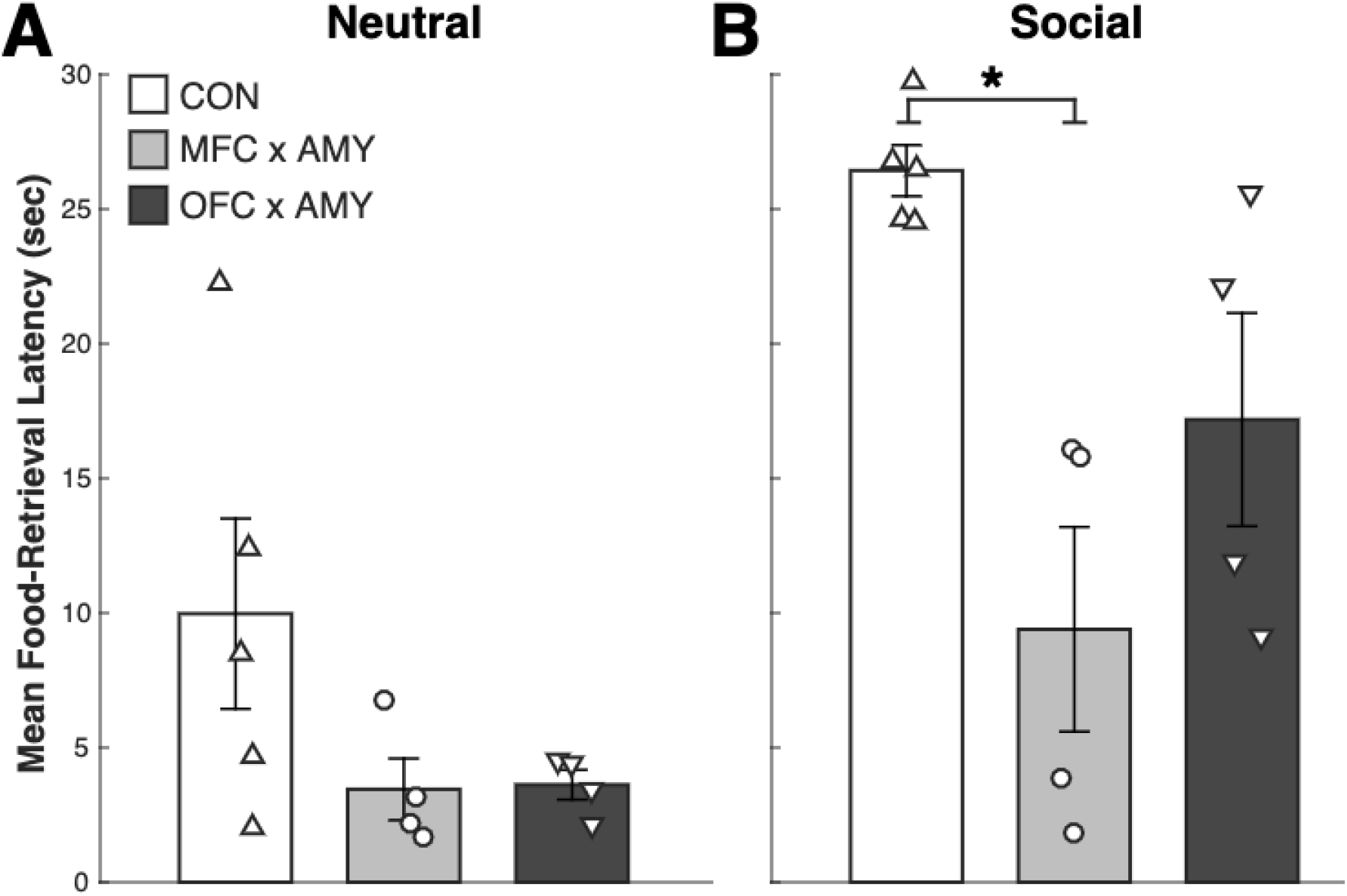
Social valuation task. **A**, Mean food-retrieval latencies when monkeys viewed videos in the neutral category. **B**, Mean food-retrieval latencies when monkeys viewed videos in the social category. Error bars represent the SEM. Groups are plotted on the X-axis and mean food-retrieval latencies (in seconds) are plotted on the Y-axis. * < 0.05.

We then ran a Kruskal-Wallis test for each category separately to test for a main effect of group. We found a significant effect of group for the social condition (*χ*^2^ = 7.78; *p* = 0.02; **Fig. 2B**) but not the neutral condition (*χ*^2^ = 3.22; *p* = 0.20; **Fig. 2A**). Specifically, we found that the MFC x AMY group was significantly faster to reach for food rewards than the control subjects in the presence of the social videos (unpaired Mann-Whitney *U* test: *W* = 20, *p* = 0.02) but not the neutral stimuli (unpaired Mann-Whitney *U* test: *W* = 16, *p* = 0.19). The OFC x AMY group did not differ from either group in the social condition (unpaired Mann-Whitney *U* test for OFC x AMY vs. MFC x AMY: *W* = 12, *p* = 0.34; OFC x AMY vs. CON: *W* = 18, *p* = 0.06, trending towards significance) or the neutral condition (unpaired Mann-Whitney *U* test for OFC x AMY vs. MFC x AMY: *W* = 10, *p* = 0.67; OFC x AMY vs. CON: *W* = 16, *p* = 0.19).

Although these data suggest that a MFC x AMY circuit is necessary for social valuation, to validate this interpretation, we attempted to rule out the possibility that subjects with MFC x AMY lesions were simply ignoring the stimuli, which might indicate a global disruption of visual attention. All groups spent more time viewing the social videos (**Fig. 3B**) compared to the neutral stimuli (**Fig. 3A**), indicated by a significant main effect of stimulus category for viewing durations (*χ*^2^ = 14.21; *p* = 0.0002). Importantly, we found no significant main effect of group on the duration of time spent looking at the social videos (*χ*^2^ = 0.10; *p* = 0.95), suggesting intact visual attention across groups.

**Figure 3.**
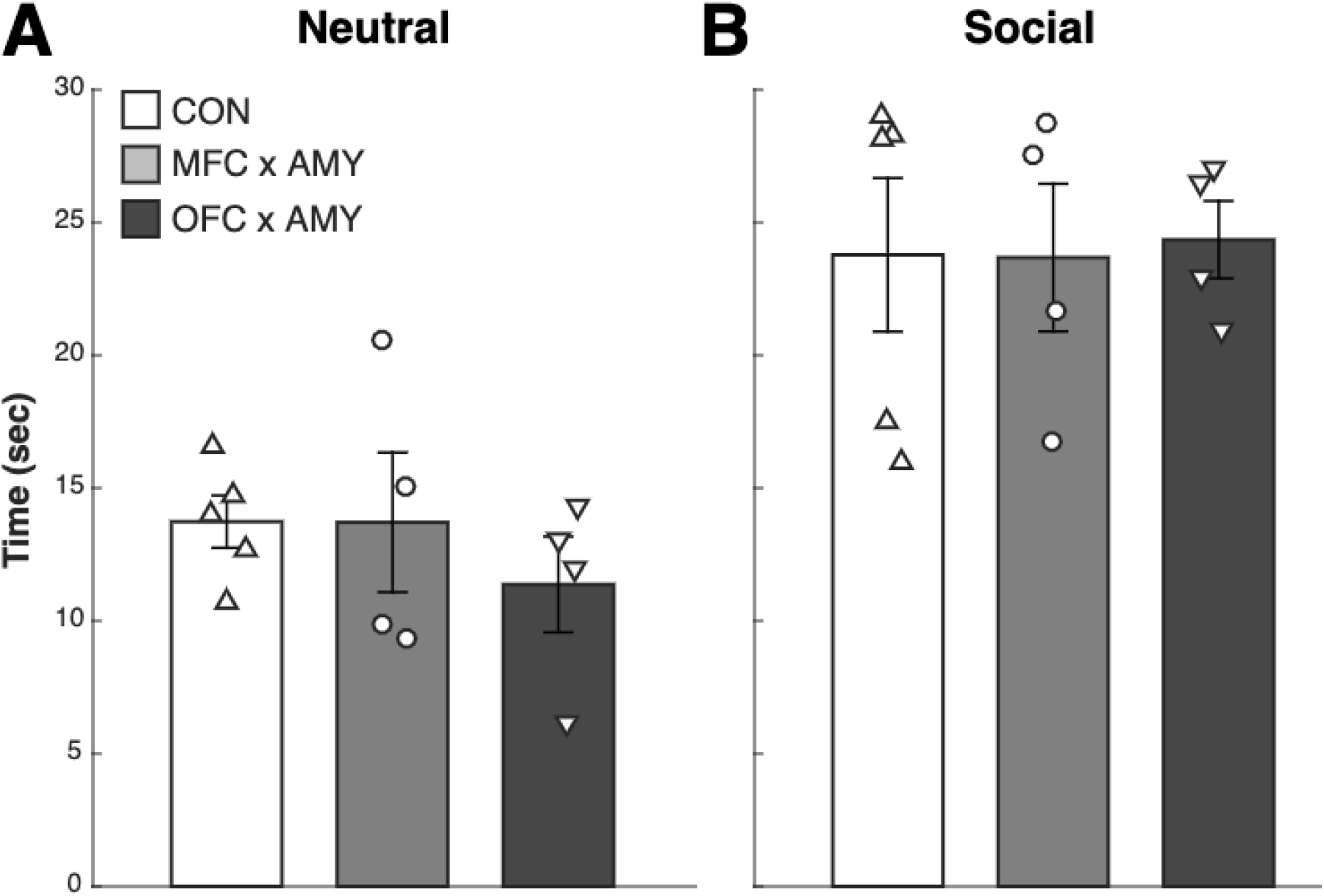
Social valuation task. **A**, Mean time spent viewing the videos in the neutral category. **B**, Mean time spent viewing videos in the social category. Error bars represent the SEM. Groups are plotted on the X-axis and mean viewing times (in seconds) are plotted on the Y-axis.

Finally, we predicted that both operated groups would show reduced food-retrieval latencies to emotionally-relevant stimuli given observed reductions in affective reactivity to predator stimuli following either unilateral (Izquierdo and Murray, 2004) or bilateral amygdala damage (Meunier et al., 1999; Izquierdo et al., 2005; Chudasama et al., 2009; Machado et al., 2009; Bliss-Moreau et al., 2011; Medina et al., 2020). However, contrary to this prediction, there was no main effect of group for food-retrieval latencies in the presence of the predator videos (*χ*^2^ = 3.2; *p* = 0.20), nor did we observe a significant effect of group for the total video viewing time (*χ*^2^ = 0.30; *p* = 0.86).

### Reinforcer Devaluation

We calculated a proportion shifted score for each subject and each test, with a higher proportion shifted score indicating a greater shift away from the object associated with the sated food. We found a significant effect of group on the proportion shifted score for object choices (*χ*^2^ = 6.50; *p* = 0.04; **Fig. 4A**). Whereas the MFC x AMY group did not differ from controls (unpaired Mann-Whitney *U* test: *W* = 12.5, *p* = 0.62), the OFC x AMY group showed significantly smaller proportion shifted scores for object choices compared to the control group (unpaired Mann-Whitney *U* test: *W* = 20, *p* = 0.02), but not compared to the MFC x AMY group (unpaired Mann-Whitney *U* test: *W* = 2, *p* = 0.11). As predicted, groups did not differ with respect to proportion shifted scores when presented with the actual foods for choice following selective satiation (*χ*^2^ = 3.66; *p* = 0.16; **Fig. 4B**). Therefore, all monkeys were able to identify foods by vision and had intact satiety mechanisms. Only monkeys with OFC x AMY lesions, however, were unable to link objects with the current value of the associated foods.

**Figure 4.**
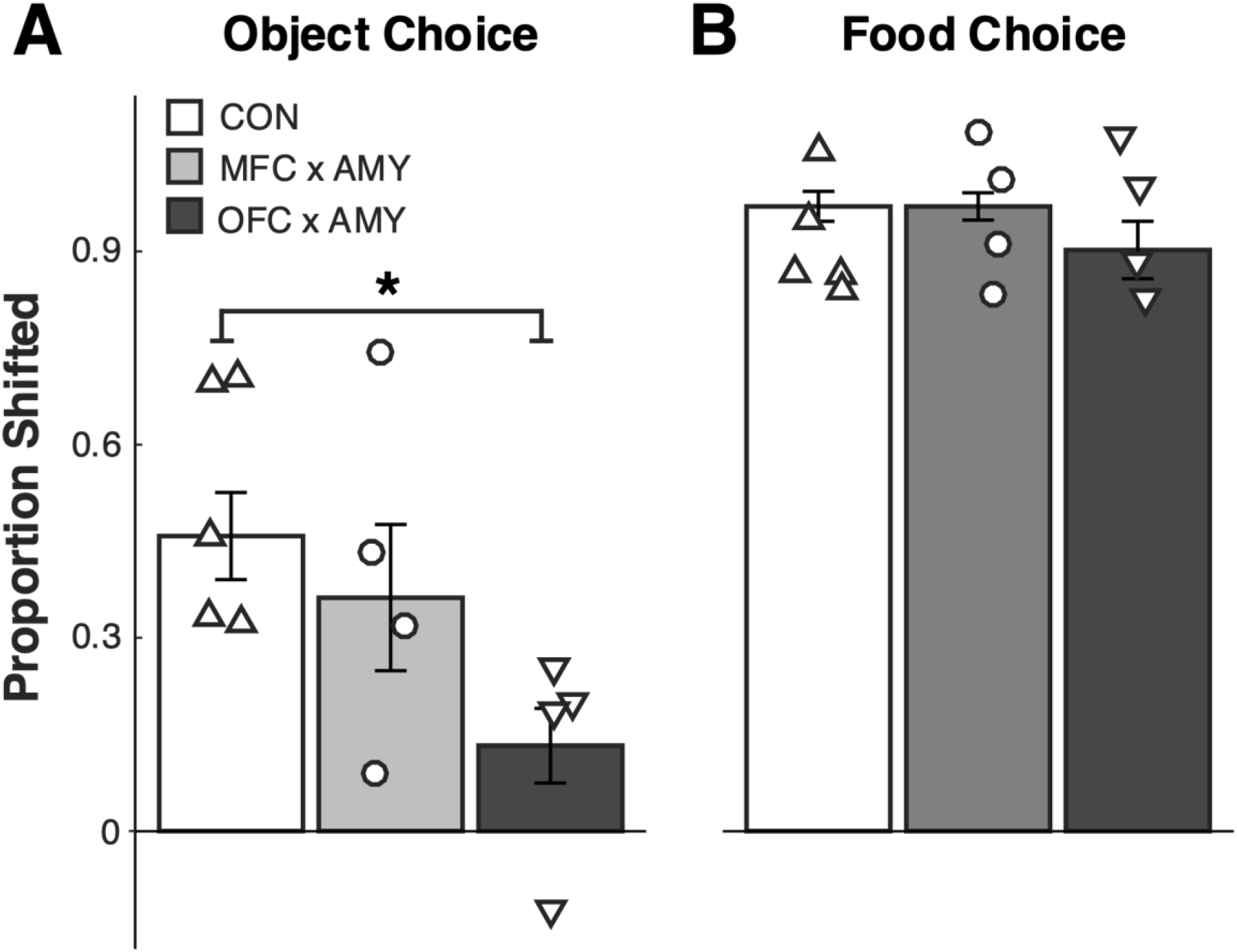
Reinforcer devaluation task. **A**, Mean proportion shifted scores for object choices. **B**, Mean proportion shifted scores for food choices. Groups are plotted on the X-axis and proportion shifted scores are plotted on the Y-axis. Error bars represent the SEM. * < 0.05.

### Double Dissociation of Functional Interactions in Prefrontal-Amygdala Circuits

Experiments 1 and 2, taken together, show that MFC interaction with the amygdala is necessary for the expression of social value, whereas OFC interaction with the amygdala is essential for updating information about object and/or food value. To demonstrate that these circuits are dissociable in terms of their designated functions, we directly compared lesion group performances on both tasks. We normalized the food-retrieval latencies from the social valuation task (social condition - neutral condition) and the average proportion shifted scores from the object choice tests. We ran a 2×2 ANOVA and found a significant group by task interaction, (*F* = 5.65; *p* = 0.03; **Fig. 5**), indicating that these circuits make selective contributions to social cognition and value-based decision making.

**Figure 5.**
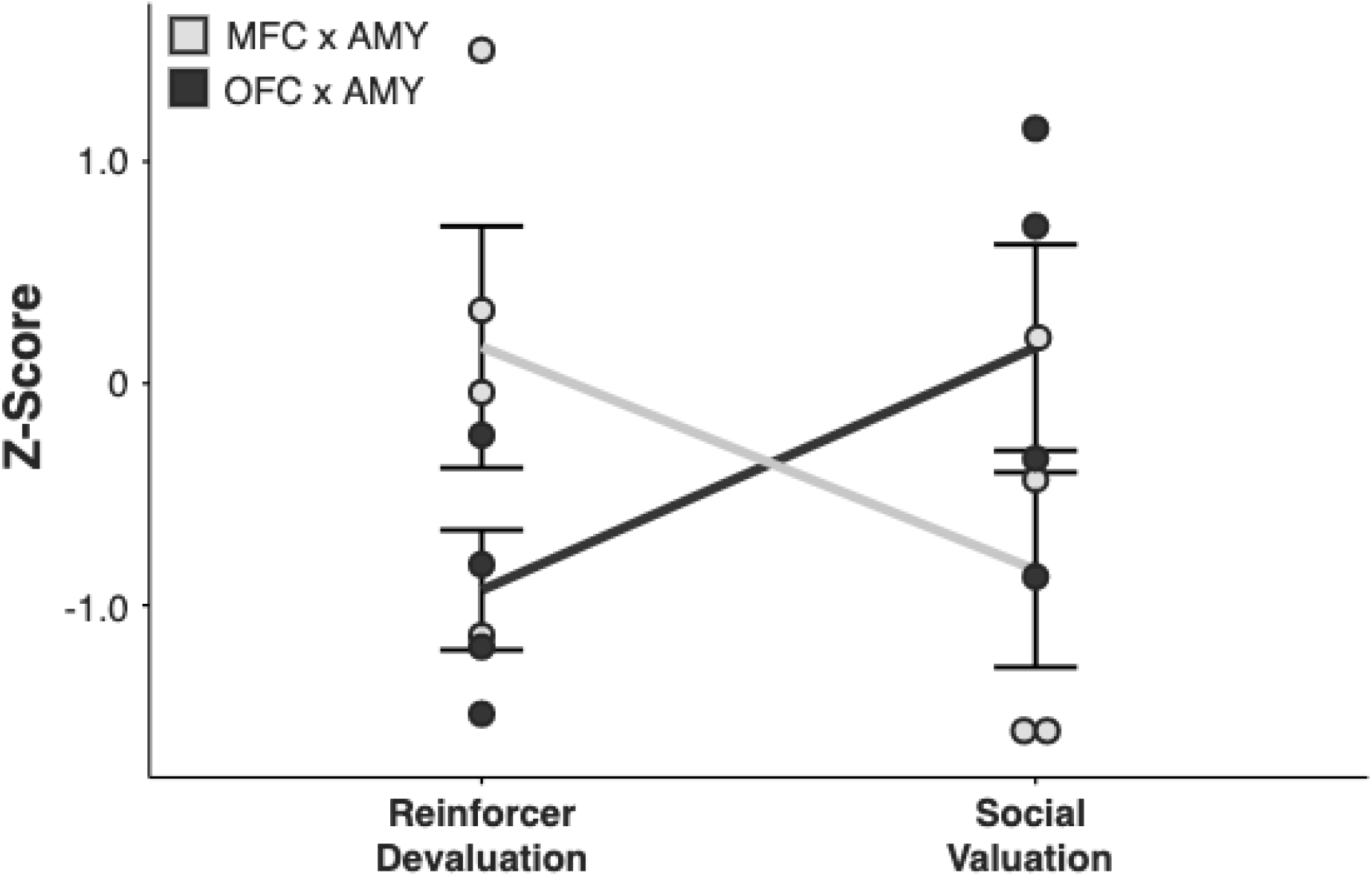
Double dissociation between groups on social valuation task performance versus reinforcer devaluation task performance. The X-axis displays each task and Y-axis displays normalized *z*-scores. Data for each group are plotted in a separate line. Error bars represent the SEM.

## Discussion

In support of our hypothesis, we showed that MFC interaction with the amygdala is necessary for the expression of social value, whereas OFC interaction with the amygdala is essential for linking objects with current food value. In a direct comparison of task performances, we demonstrated that these prefrontal-amygdala circuits make selective and dissociable contributions to behavior.

### Mechanisms of MFC-amygdala interaction during social cognition

A wealth of evidence indicates that the MFC supports social cognition. For example, MFC damage in macaques yields reduced sensitivity to social information (Hadland et al., 2003; Rudebeck et al., 2006), and single unit recordings reveal a population of neurons in the MFC that encodes information about rewards for “other” vs. “self” (Chang et al., 2013). Decades of human neuroimaging work has resulted in the characterization of a so-called “social brain network,” that includes the amygdala, MFC, and superior temporal sulcus/temporo-parietal junction (Blakemore, 2008; Apps et al., 2016; Noonan et al., 2017; Burgos-Robles et al., 2019; Lockwood et al., 2020; Gangopadhyay et al., 2021). Recently, it was found that monkeys with bilateral, excitotoxic MFC lesions, unlike controls, failed to develop prosocial tendencies in a reward allocation learning task involving the selection of visual cues that predicted reward delivery to the self, another monkey, or no one (Basile et al., 2020). In a recent multi-site neural recording study in monkeys using a variant of the same task, oscillatory interactions between the MFC (specifically the ACC, including areas 32, 24b, and 24a) and amygdala were enhanced when monkeys chose to give juice to another monkey, but suppressed when a monkey acted antisocially (Dal Monte et al., 2020). Although our experiment did not require associative learning in a social context, these and our results warrant future studies to examine whether learned cue-outcome associations from socially-derived information are disrupted following transient or chronic interruption of MFC-amygdala activity.

Although at least two studies have found no effects of OFC damage on social cognition in macaques (Rudebeck et al., 2006; Noonan et al., 2010b), there is some evidence in support of OFC involvement in social cognition, as well (Watson and Platt, 2012; Goursaud and Bachevalier, 2020). One line of evidence comes from the discovery of face patches in the lateral orbital sulcus of the OFC (Tsao et al., 2008; Troiani et al., 2016). Recording studies suggest that these patches contain face-selective neurons that categorize social information from faces (Barat et al., 2018). In our own data, the OFC x AMY group shows a trend-level difference from controls on their food-retrieval latencies in the presence of the social videos. This raises the possibility that the division of labor between MFC and OFC in social cognition may not be so clear-cut, and that perhaps an intra-PFC network involving subregions of MFC and OFC encodes information that is relevant to social value and learning from social contexts. Another possibility, not mutually exclusive with the first, is that unilateral amygdala damage produces a mild disruption of social evaluation.

The amygdala has ubiquitously been implicated in social cognition (Adolphs et al., 1998; Amaral, 2002; Adolphs, 2010). Recently, Grabenhorst et al. (2019) reported that a population of neurons in the macaque amygdala simulate a conspecific’s decisions prior to their choice during an observational learning task involving juice rewards. An intracranial recording study in humans also presents the possibility that there are two populations of neurons in the amygdala which encode cue-outcome associations from experienced (self) and observed (other) outcomes (Aquino et al., 2020). Another recent study in macaques found a population of amygdala neurons that code for both juice reward value and conspecific social rank, whereas the OFC and ACC coded for reward values but not hierarchical rank (Munuera et al., 2018). Taken together, and consistent with a role for MFC-amygdala interactions in social cognition, these data suggest that the amygdala may be a site where stimulus-outcome value associations and social information are integrated.

### Mechanisms of OFC-amygdala interactions during value updating

Individually, the OFC (Izquierdo et al., 2004; Machado and Bachevalier, 2007; Noonan et al., 2010b; Rudebeck et al., 2013b; Rudebeck et al., 2017a) and the amygdala (Malkova et al., 1997; Izquierdo and Murray, 2007), but not the MFC (Chudasama et al., 2013; Rhodes and Murray, 2013), are necessary for nonsocial, value-based decisions in macaques. The present study confirms the results of an earlier study examining OFC-amygdala disconnection (Baxter et al., 2000) and extends those findings in two ways. First, the present study, unlike the earlier one, used selective, excitotoxic lesions of OFC. Thus, the results confirm an essential role in decision making for amygdala projections to or from neurons in OFC. Second, the present study, unlike the earlier one, did not include a section of the forebrain commissures as part of the surgical disconnection. That the impairments in value updating follow from crossed surgical disconnection with the forebrain commissures intact indicates that the behavior depends on intrahemispheric interactions of the amygdala and OFC. In addition, previous work in macaques demonstrates a critical role for amygdala inputs to OFC but not MFC in sustaining reward value coding for well learned stimulus-reward amounts (Rudebeck et al., 2013a) and in acquiring value coding in terms of both the information signaled and the proportion of neurons encoding stimulus–reward amounts during learning (Rudebeck et al., 2017b).

The possible mechanism by which OFC and amygdala interactions subserve value updating is also starting to be unraveled. In macaques, pharmacological inactivation of posterior OFC (Murray et al., 2015) or the basolateral amygdala (Wellman et al., 2005) during but not after selective satiation impairs adaptive shifts in object choices on the devaluation task. Together, these studies imply that the OFC critically interacts with the amygdala to register changes in food value during the selective satiation procedure, and that this updated information is used to guide subsequent goal-directed choices. A mechanistic account has been proposed wherein the basolateral amygdala is involved in updating stimulus-outcome associations and the OFC maintains representations of past and present associations that are retrieved by the basolateral amygdala in the updating process (Sharpe and Schoenbaum, 2016; Lichtenberg et al., 2017). Thus, communication between the amygdala and the OFC appears to be critical for accurate value representations and cue-outcome associations to allay maladaptive choices.

### Limitations and future directions

There are a number of limitations that may be addressed by future work. First, our social valuation task constitutes just one of many approaches for probing social cognition. Because only four discrete stimuli of conspecifics were utilized, we were unable to reveal potentially interesting differences in reactivity among different classes of social stimuli (e.g., submissive vs. dominant, female vs. male). The small number of subjects in each group also precluded our ability to determine whether dominance status of the monkeys changed following the lesions and whether dominance status at the time of testing was a contributing factor in video evaluation. Additionally, our measure of gaze duration for the stimuli was a crude estimation of video fixation duration. A follow-up study employing eye-tracking measures for multiple social stimulus categories would isolate the effects of attention from social valuation by allowing the detection of subtle differences in gaze-fixation patterns between social and nonsocial stimuli among lesion and control groups, if any, as in Deaner et al. (2005).

Another limitation in our study is the use of aspiration lesions to target the MFC. We cannot rule out the possibility that the behavioral effects of the MFC lesions were due to unintended damage to fibers of passage. Also, the area of the MFC targeted in our study may not have covered the full extent of the medial wall within the boundaries of the ACC thought to be involved in social cognition (Apps et al., 2016). Importantly, however, the MFC cortex lesion in our study included the gyrus of the ACC, which was found to be critical in supporting prosocial behaviors, relative to the sulcus of the ACC which has not been implicated in these behaviors (Rudebeck et al., 2006; Chang et al., 2013).

## Conclusion

In this study, we show that the MFC and amygdala must interact to support the ability to ascribe value to others. Dysfunction in this circuitry may underlie social affiliation deficits – that is, marked reductions in an interest in and/or empathy for others – observed in psychopathy (Viding and McCrory, 2019) and autism spectrum disorder (Thakkar et al., 2008; DeMayo et al., 2019). We further show that the OFC and amygdala form a functional circuit that is essential for sustaining adaptive stimulus choice behaviors following changes in the incentive value of food rewards. Dysregulation in this circuit may underlie disorders in which compulsive and perseverative behaviors are observed, such as substance use disorder (Koob and Volkow, 2016; Bechara et al., 2019) and obsessive-compulsive disorder (Gillan and Robbins, 2014). The framework presented in the current study regarding the separable functions of these cortico-amygdala circuits will require additional testing to determine the anatomical specificity, molecular substrates, and temporal dynamics that give rise to these complex behaviors.

## Acknowledgements

This work was supported by the Intramural Research Program of the National Institute of Mental Health (EAM, ZIAMH002887). MSP was supported by National Institutes of Health Postdoctoral Fellowship Center for Compulsive Behavior Postdoctoral Fellowship. We thank Dawn Lundgren, Emily Moylan, and Emily Fiuzat for assistance with data collection; Richard Saunders and Emily Moylan for help performing surgery; and Ben Jung, Jakob Seidlitz, and Adam Messinger for assistance with the lesion assessments. We also thank the staff of the Nuclear Magnetic Resonance Facility, National Institute of Neurological Disorders and Stroke, and the Laboratory of Diagnostic and Radiology Research. Current author affiliations: MSP (Sarah Lawrence College, 1 Mead Way, Bronxville, NY 10708); NKC (Temple University, 1701 N Broad St, Philadelphia, PA 19122); SEVR (Division of Program Coordination, Planning, and Strategic Initiatives, National Institutes of Health, Bethesda, MD 20892).

